# A subtle modification of modafinil-based DAT inhibitors changes conformational preference

**DOI:** 10.1101/2024.03.06.583808

**Authors:** Kuo Hao Lee, Gisela A. Camacho-Hernandez, Amy Hauck Newman, Lei Shi

## Abstract

Modafinil analogs with either a sulfoxide or sulfide moiety have improved binding affinities at the human dopamine transporter (hDAT) compared to modafinil, with lead sulfoxide-substituted analogs showing characteristics of atypical inhibition (e.g., JJC8-091). Interestingly, the only distinction between sulfoxide and sulfide substitution is the presence of one additional oxygen atom. To elucidate why such a subtle difference in ligand structure can result in different typical or atypical profiles, we investigated two pairs of analogs. Our quantum mechanical calculations revealed a more negatively charged distribution of electrostatic potential surface of the sulfoxide substitution. Using molecular dynamics simulations, we demonstrated that sulfoxide-substituted modafinil analogs have a propensity to attract more water into the binding pocket. They also exhibited a tendency to dissociate from Asp79 and form a new interaction with Asp421, consequently promoting an inward-facing conformation of DAT. In contrast, sulfide-substituted analogs did not display these effects. These findings deepen our understanding of the functionally relevant conformational spectrum of DAT.

## Introduction

The primary function of the dopamine transporter (DAT) is to regulate dopamine levels in the synaptic cleft by transporting the released dopamine back into the presynaptic dopaminergic neurons, thereby terminating dopamine neurotransmission.^1, 2^ Disruption of dopamine uptake and signaling has been associated with various psychiatric disorders, including schizophrenia, Parkinson’s disease, bipolar disorder, depression, and attention deficit hyperactivity disorder (ADHD).^3–5^ Cocaine binds to all three monoamine transporters, but its psychostimulant actions have been primarily attributed to inhibition of DA reuptake via DAT.^6–8^ Consequently, this inhibition results in a rapid accumulation of dopamine in the synaptic cleft that is widely regarded as the primary cause of the reinforcing effects of cocaine that can lead to addiction, in humans.^9^ Indeed it has been hypothesized that all compounds that inhibit DAT would induce behaviors similar to those caused by cocaine.^10–12^ However, many studies have demonstrated that certain DAT inhibitors, including benztropine, modafinil and their analogs, exhibit diminished rewarding effects, in humans and experimental animals, and thus reduced liability for misuse and addiction.^13–16^ The term “atypical DAT inhibitors” has been designated to refer to such compounds, whereas substances such as cocaine and cocaine-like compounds are categorized as “typical DAT inhibitors”.^13^

DAT belongs to the Neurotransmitter:sodium symporter family, the members of which translocate substrates across the cellular membrane by utilizing the energy stored in the transmembrane Na^+^ gradient.^8^ In DAT, there are two sodium binding sites, each playing distinct roles. The first sodium ion (Na1) is closely associated with substrate binding, while the bound second sodium ion (Na2) is crucial in stabilizing the outward-facing state.^17–20^ In the transport process, DAT undergoes conformational transitions by alternating between outward-facing (including outward-open and outward-occluded) and inward-facing (including inward-open and inward-occluded) states.^21, 22^ X-ray crystallography has captured the outward-open, outward-occluded, and inward-open conformations of the Leucine transporter (LeuT), a bacterial homolog of DAT.^23–26^ Various substrates and inhibitors have been co-crystallized with *Drosophila melanogaster* DAT (dDAT), which has a sequence identity of more than 50% with human DAT (hDAT).^27, 28^ The dDAT structures were all in outward-facing conformations.

Based on the crystal structures of LeuT and dDAT, computational modeling and simulations have been instrumental in gaining insights into the molecular mechanisms of ligand binding and conformational dynamics in DAT.^22, 29–36^ It is essential to identify and characterize the specific conformations that are differentially stabilized by typical and atypical inhibitors to understand how these inhibitors function. Specifically, the Y335A DAT mutation, which promotes the inward-open conformation of DAT, was observed to have a lesser impact on the binding of atypical DAT inhibitors compared to cocaine-like inhibitors.^37^

The mechanisms through which atypical inhibitors affect the DAT and the resulting conformational rearrangements in the protein have been explored through molecular dynamics (MD) simulations. Specifically, our previous study indicated that JHW007, a benztropine analog, preferred an inward-occluded DAT conformation.^37, 38^ Moreover, JJC8-091, a modafinil analog that does not display any cocaine-like behaviors in rodents, favored a more occluded DAT conformation in comparison to its derivative, JJC8-088, a modafinil analog that is fully cocaine-like in rats and nonhuman primates.^39, 40^ MD simulations have proven valuable in unveiling these conformational changes induced by different inhibitors, providing insights into the molecular intricacies associated with the transition from atypical to typical inhibitors and vice versa.

The atypical effects of modafinil and its derivatives have been the subject of studies exploring the correlation between these effects and conformational changes in DAT.^41, 42^ For example, JJC8-091, is not self-administered in experimental animals and is capable of preventing cocaine-induced reinstatement of drug-seeking behavior.^40^ As a result, it has been further developed as a promising candidate for the treatment of psychostimulant addiction, although recent studies in nonhuman primates were less convincing than in rats.^16^

In this study, we carried out quantum mechanical calculations and extensive MD simulations to comparatively characterize the interactions between hDAT and two pairs of modafinil analogs, JJC8-091 and RDS04-010 with the sulfoxide substitution and their sulfide analogs JJC8-089 and RDS03-094, respectively. The analysis of our MD simulation results demonstrated that the subtle difference between sulfoxide versus sulfide impact the conformational equilibrium of hDAT and is predicted to result in divergent behavioral profiles.

## Results and Discussion

### The sulfoxide and sulfide modafinil analogs have distinct physical-chemical properties

The only difference between RDS03-094 and RDS04-010 (referred to as RDS pairs in this study) is the substitution of sulfoxide (-S(=O)-) verses sulfide (-S-). Similarly, JJC8-091 and JJC8-089 (referred to as JJC pairs) also have the same difference. The results of our binding assay showed that the sulfoxide analogs in both pairs have ∼30 fold lower affinities at hDAT compared to the corresponding sulfide analogs (**Table 1**). Compared to the JJC pairs, the RDS pairs have a 2,6-dimethyl-substitution on the piperazine ring, which had no significant effect on DAT binding affinities (**Table 1**). To compare the impact of the sulfoxide and sulfide substitution on the compound conformation and electrostatic properties, we first carried out conformational search of each compound to identify their conformers with the lowest energy. Although the lowest-energy conformers of these two pairs of analogs share significant similarity in their extended conformations, they demonstrate noticeable difference near their aligned sulfoxide and sulfide moieties.

**Table 1.**
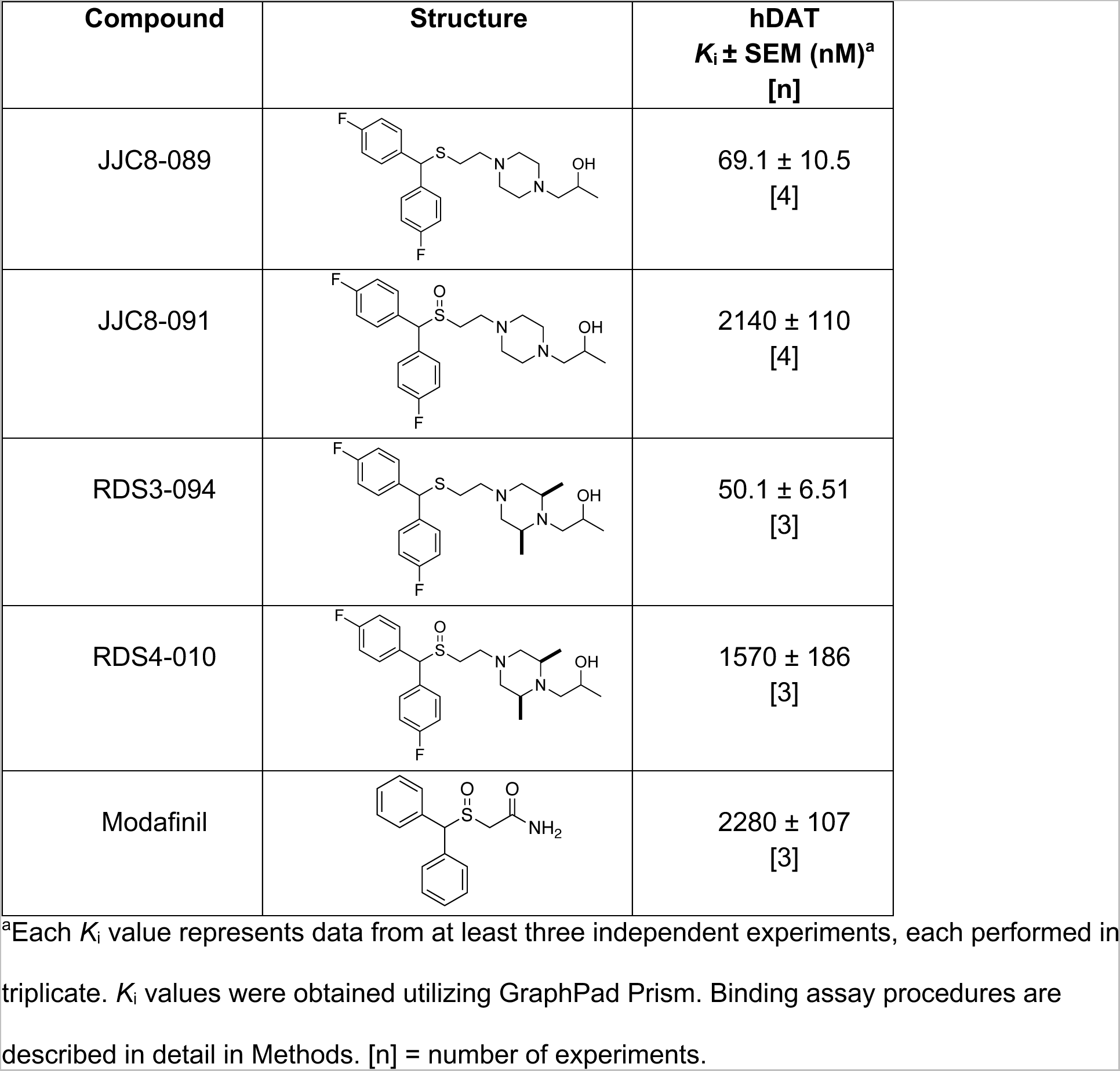
Binding affinities of the modafinil analogs.

| Compound | Structure | hDAT<br>$K_i \pm \text{SEM}$ (nM)<sup>a</sup><br>[n] |
| --- | --- | --- |
| JJC8-089 | | $69.1 \pm 10.5$<br>[4] |
| JJC8-091 | | $2140 \pm 110$<br>[4] |
| RDS3-094 | | $50.1 \pm 6.51$<br>[3] |
| RDS4-010 | | $1570 \pm 186$<br>[3] |
| Modafinil | | $2280 \pm 107$<br>[3] |

<sup>a</sup>Each $K_i$ value represents data from at least three independent experiments, each performed in triplicate. $K_i$ values were obtained utilizing GraphPad Prism. Binding assay procedures are described in detail in Methods. [n] = number of experiments.

Using these lowest-energy conformers as the inputs, we then further optimized geometry of these conformers and characterized their electrostatic potential surfaces (EPS) with quantum mechanical (QM) calculations. Using the RDS pair as an example, the lowest energy conformers show a relatively extended conformation similar to those resulting from the conformational search (**Fig. 1C, D**). As expected, the most prominent difference on their EPS is the negative partial charge distributed around the oxygen atom of the sulfoxide (RDS04-010), which is absent in the sulfide (RDS03-094) (**Fig. 1A, B**). These different charge distributions lead to varied orientations between the bisphenyl and 2,6-dimethylpiperazine moieties of these two compounds. Based on the optimized geometries of RDS04-010 and RDS03-094 from the QM optimization, the dihedral angle between those two moieties can vary from −35.8° to −46.0° due to the presence of sulfoxide substitution (**Fig. 1C, D**). The differences in both charge distribution and ligand conformation may result in varied propensity to form either ligand-water or ligand-protein interactions (see below).

**Figure 1.**
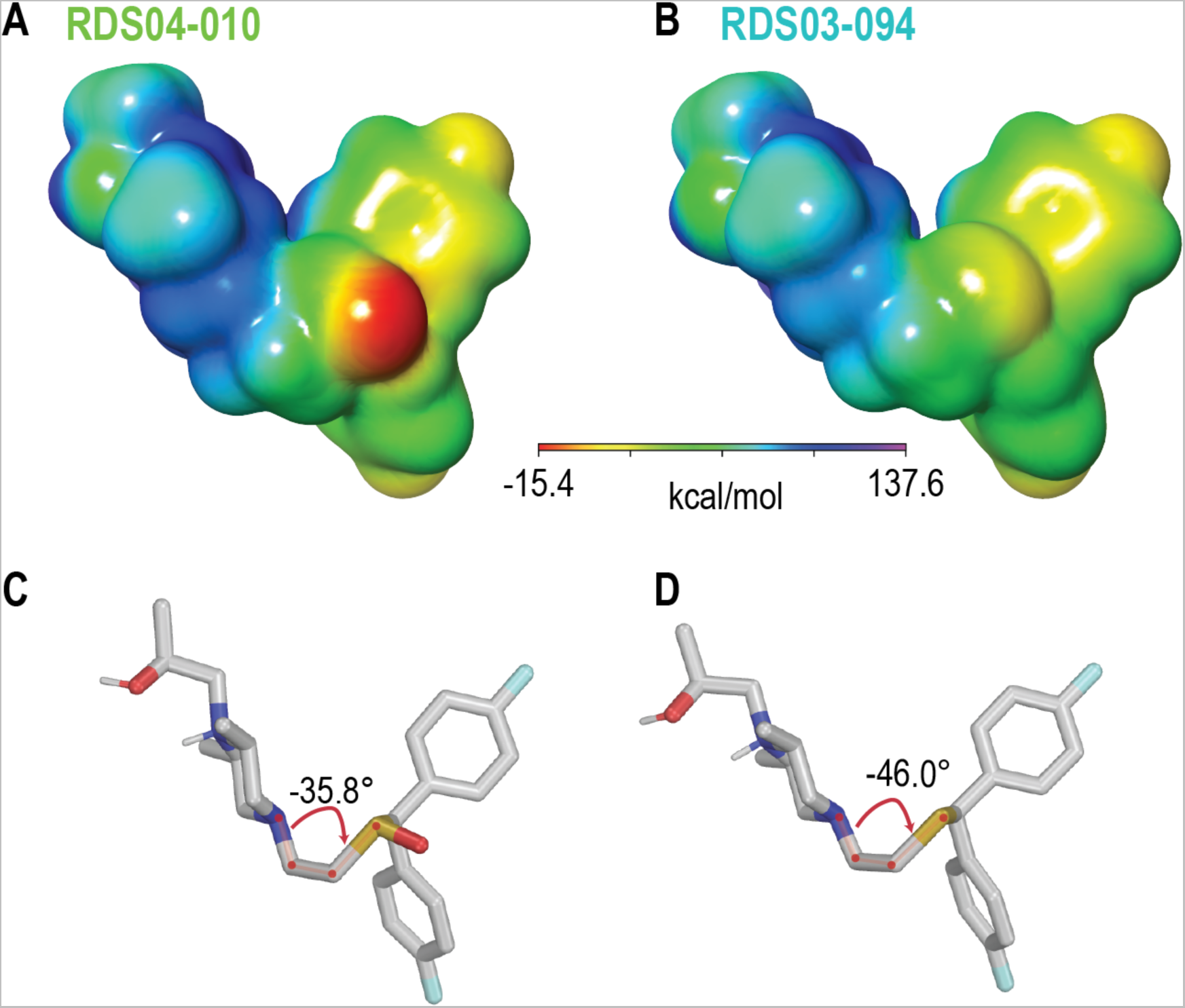
The addition of the sulfoxide moiety significantly alters the physical-chemical properties of the ligand. The calculated electrostatic potential surfaces for the RDS04-010 (**A**) and RDS03-094 (**B**) shows that the sulfoxide introduces partial negative charge (red) near the bisphenyl moiety. As a result, the dihedral angle between the bisphenyl and dimethylpiperazine moieties has > 10° difference between RDS04-10 (**C**) and RDS03-094 (**D**).

### The sulfoxide and sulfide containing-ligands have different binding poses at hDAT

To evaluate the differential impact of the sulfoxide versus sulfide substitution on the binding of the ligands with the same scaffold at hDAT, we docked the ligands into a hDAT model in an outward-facing conformation, immersed the hDAT/ligand complexes individually in explicit lipid bilayer/water environment, and performed prolonged molecular dynamics (MD) simulations (see Methods, **Table S1**, and **Fig. 2A**).

**Figure 2.**
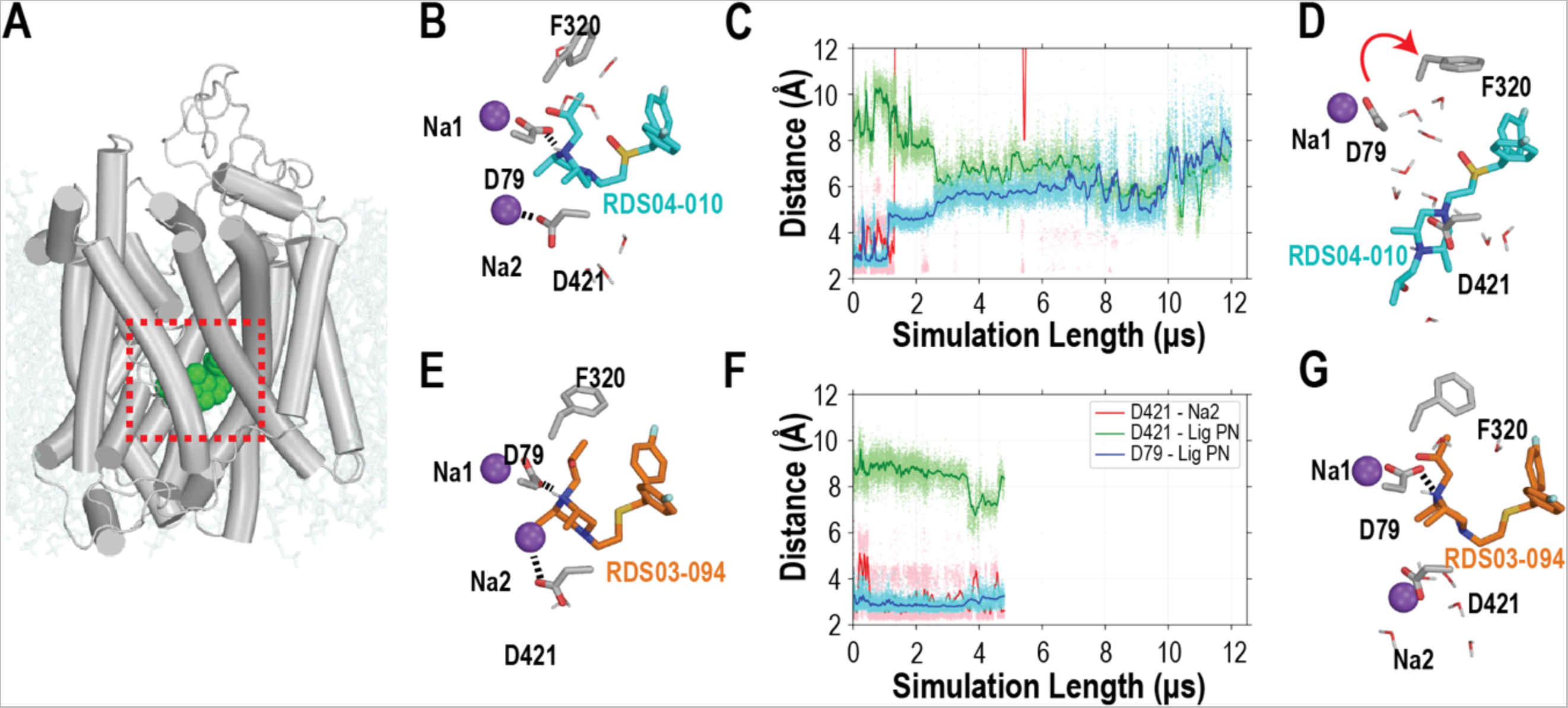
The bound of RDS04-010 cannot stably interact with Asp79 resulting in the dissociation of Na2. (**A**) hDAT in complex with RDS04-010 is embedded in lipid bilayer (pale cyan) with the central ligand binding pocket indicated by a red box. The initial binding poses of RDS04-010 and RDS03-094 from docking are shown in **Fig. 2B** and **E** respectively, and the representative snapshots of their equilibrated binding poses are shown in **Fig. 2D** and **G**, respectively. During the MD simulations, as shown in panels **B**-**D** from a representative trajectory, RDS04-010 gradually loses its interaction with Asp79, which is present in the starting model (**B**), and gets close to Asp421 at the end of the trajectory (**D**); the evolution of the distances between the pyramidal nitrogen of the ligand and these two residues are shown in panel **C**. The dissociation of Na2 from its binding site, which is demonstrated by its elongated distance to Asp421, is prerequisite for the transition to happen. In comparison, such a transition does not happen in the simulations of hDAT bound with RDS03-094 (**E**-**G**). The sodium ions (Na1 and Na2) are shown in purple. For simplicity, the stably bound chloride ion is not illustrated. Note that the distance between ligand pyramid nitrogen and Asp421 or Asp79 was the minimum distance between pyramidal nitrogen and the sidechain oxygen atoms. Similarly, the distance between Asp421 and Na2 was the minimum distance between Na2 and the sidechain oxygen atoms of Asp.

The inspection of the resulting hDAT models from the equilibrated simulations indicate that these two substitutions in both the RDS and JJC pairs result in similarly different interactions within the binding pocket. While these four ligands commonly interact with 26 residues in TMs1, 3, 6, 8, and 10 (**Table S2**), overall, the ligand binding pose tends to be more extended for the ligands with the sulfoxide compared to those with the sulfide. Consequently, residues Gly425 of TM8 interact more frequently with the sulfoxide-containing ligands, while Ala81 of TM1, Asn157 of TM3 and Ala479 of TM10 interact more frequently with the sulfide-containing ligands. In addition, a direct ionic interaction was formed persistently between the pyramidal nitrogen of the ligands (L_N_) and the negatively charged Asp79, in hDAT/RDS03-094 and hDAT/JJC8-089 throughout the simulations (**Figs. 2E-G and S1D-F1**), while this interaction was gradually lost during the hDAT/RDS04-010 and hDAT/JJC8-091 simulations (**Figs. 2B-D and S1A-C**). The divergence at this critical interaction is likely reflected in the higher affinities of RDS03-094 and JJC8-089 compared to RDS04-010 and JJC8-091, respectively (**Table 1**).

In addition, the hDAT/RDS04-010 and hDAT/JJC8-091 simulations showed a common tendency for the more intracellular bound Na^+^ ion near the ligand binding pocket (Na2) to escape to the intracellular milieu (**Fig. 2B, D, E, and G**), which exposed a negatively charged Na2 coordinating residue, Asp421, to potentially attract and interact with the ligands. Note that such a Na2-escaping tendency has been previously associated with the transition to the inward-facing conformations in hDAT and homologous transporters.^17, 19, 30, 31^

To further understand the coordination among these structural elements, we compared the evolution of the distances between the L_N_ of the ligands and Asp79 and Asp421, as well as that between Na2 and Asp421 (**Fig. 2C, F**). In the hDAT/RDS04-010 simulations, in eight out of nine trajectories, Na2 was not stable and the ionic interaction between Na2 and Asp421 gradually dissociated (i.e., the distance > 5 Å). The escape of Na2 from the binding pocket resulted in an opening at the bottom of the ligand binding site (**Fig. 2**), while the L_N_ of RDS04-010 gradually dissociated with Asp79 before forming a water-mediated interaction with Asp421. In contrast, in the hDAT/RDS03-094 simulations, Na2 remains stably bound in the Na2 binding site, while the ionic interactions of Na2-Asp421 and L_N_-Asp79 persistently formed. The same coordination among these elements was also observed for the hDAT/JJC8-091 but not for hDAT/JJC8-089 simulations (**Fig. S1B, E**).

### The more extended pose of RDS04-010 is associated with the more polar nature of the sulfoxide

The conformational transition between the inward-facing and outward-facing conformations of transporter proteins could be characterized by the changes of the volumes of the intracellular and extracellular vestibules (IV and EV, respectively). Previously, we developed a protocol to quantitatively estimate these volumes by the numbers of water molecules detected in these vestibules (referred to as “water count”, see Methods).^17, 38, 43^ Using this protocol, our analysis showed that the representative frame ensemble from the hDAT/RDS03-094 condition has an average IV water count of 31.3, while that of hDAT/RDS04-010 has an average IV water count of 54.2. Therefore, the hDAT/RDS03-094 condition exhibited a notably smaller IV volume compared to the presence of the bound RDS04-010. On the extracellular side, hDAT/RDS03-094 showed a significantly higher average EV water count of 73.7, in contrast to the 62.0 observed in hDAT/RDS04-010 (**Fig. 3**). A similar trend was also evident in the JJC pair (**Fig. S2**). These findings further confirmed that RDS03-094 induced the outward-facing state in hDAT, while RDS04-010 promoted the inward-facing state.

**Figure 3.**
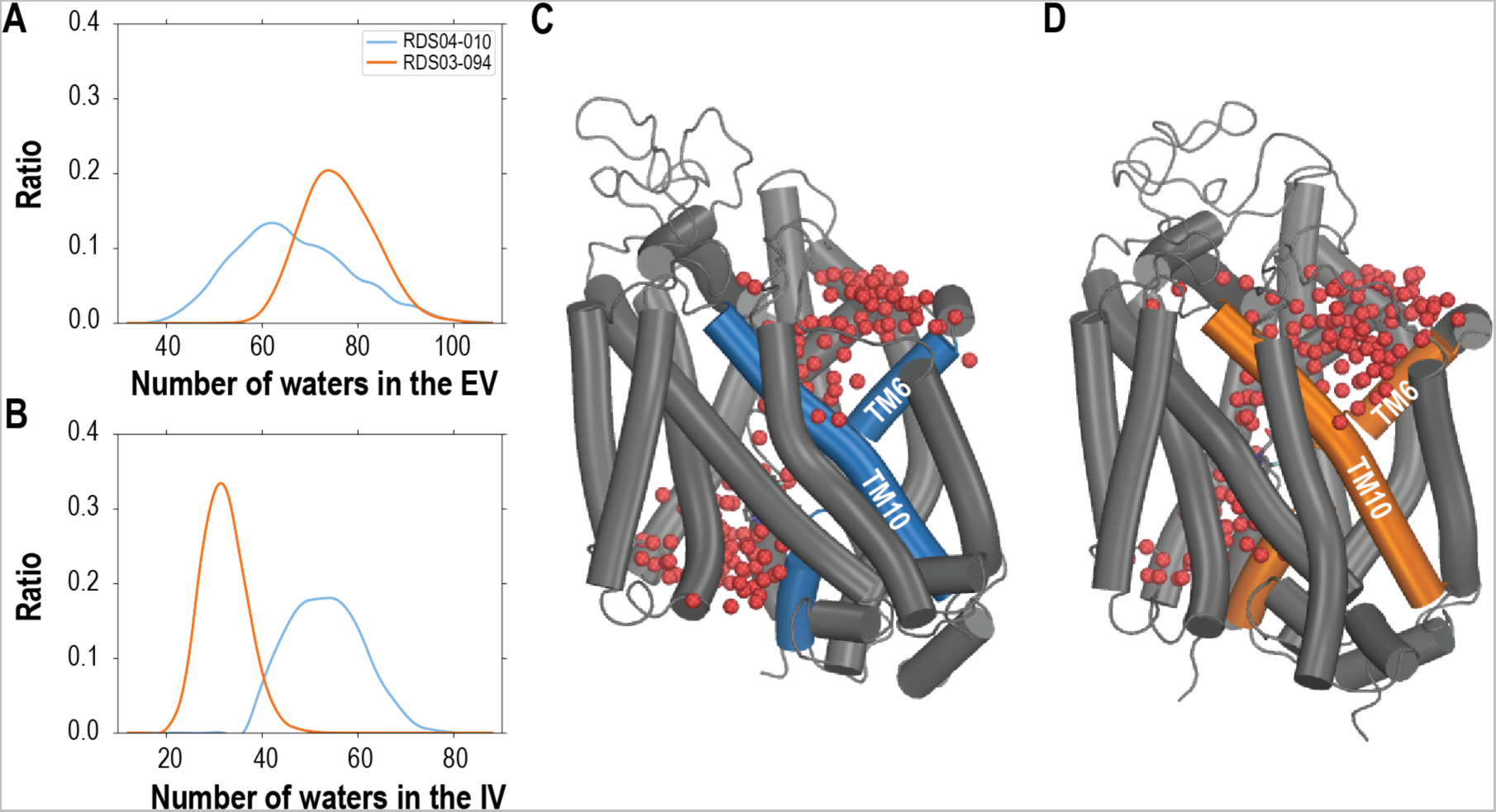
The outward-facing and inward-facing hDAT conformations stabilized by RDS04-010 and RDS03-094, respectively, show drastically different extracellular- and intracellular-vestibule volumes. We measured the volumes by counting the water molecules in the vestibules (see Methods) and found that hDAT/RDS03-094 (orange) has ∼11.7 more waters in the extracellular vestibule (**A**) and ∼22.8 fewer waters in the intracellular vestibule (**B**) than hDAT/RDS04-010 (blue). These differences are demonstrated in the hDAT/RDS04-010 (**C**) and hDAT/RDS03-094 (**D**) models with the water molecules shown in red spheres.

By extracting and comparing the frame ensemble corresponding to the inward-facing and outward-facing states, we further compared the key structural features of RDS04-010 and RDS03-094 binding in their preferred inward-facing and outward-facing hDAT conformations. We found that the distribution of the ligand end-to-end distances has the peak value of 5.6 Å for RDS03-094 in the outward-facing hDAT conformational ensemble and 7.3 Å for RDS04-010 in the inward-facing one (**Fig. S3**). This difference corresponds well to the more extended binding pose of RDS04-010, which has the sulfoxide substitution.

Furthermore, as the sulfoxide has a greater distribution of negative charge compared to the sulfide (see **Fig. 1**), more water molecules were attracted to the sulfoxide of the ligands (< 3.5 Å) in the simulations than to sulfide, as the sulfoxide but not the sulfide can form direct H-bond with these water molecules (**Fig. S4**). Thus, the presence of oxygen in the sulfoxide, not only impacts its charge distribution and overall ligand conformations, but also the tendency to be exposed to a polar environment, which ultimately contributes to the more extended ligand binding poses in hDAT (**Figs. 2 and S1**).

### The RDS04-010 stabilized hDAT conformation shows key features of inward-facing state

We then further characterized the overall conformations of the transmembrane domain of hDAT using the protein interaction analyzer (PIA),^44, 45^ which provides a quantitative assessment of conformational differences between two conditions without relying on protein superposition. To achieve this goal, we have previously defined extracellular, middle, and intracellular subsegments for each TM segment of DAT (see Methods for subsegment definitions) (**Fig. 4A**).^38, 44^

In comparing the hDAT conformation stabilized by RDS04-010 and that stabilized by RDS03-094, we found that the hDAT/RDS04-010 condition has many smaller pairwise distances among the extracellular subsegments (reddish pixels in **Fig. 4B**), while having many larger distances on the intracellular side compared to hDAT/RDS03-094 (**Fig. 4C**). Specifically, in hDAT/RDS04-010, when compared to hDAT/RDS03-094, the TM subsegments TM1e, TM6e, TM7e, TM11e, EL3, and EL4 moved closer to TM10e on the extracellular side, while TM1i swings out towards the lipid membrane with a coordinated rearrangements of TM5i, TM6i, and TM7i on the intracellular side (**Fig. 4D, E**). Thus, hDAT/RDS04-010 has clearly transitioned into an inward-facing conformation, while hDAT/RDS03-094 remained in an outward-facing conformation. A similar trend of these rearrangements was also observed between the JJC pair (**Fig. S5**).

**Figure 4.**
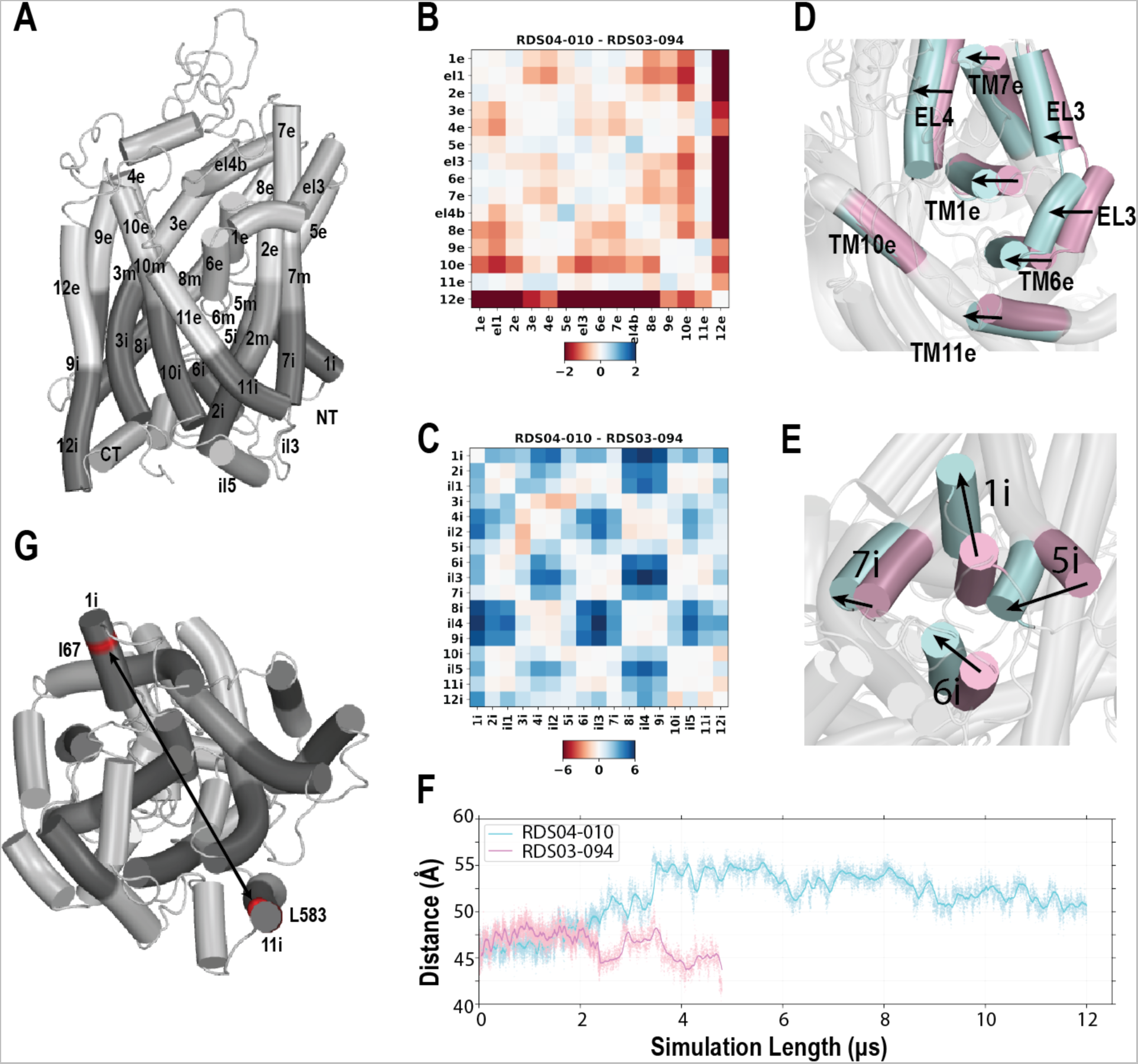
The binding of RDS04-010 induces hDAT to transition from the outward-facing to the inward-facing conformation. Panel **A** shows the division of transmembrane segments of hDAT into the extracellular (“e”), middle (“m”), and intracellular (“i”) subsegments. The differences in the distances among the subsegments shows that compared to the hDAT model bound with RDS03-094 (hDAT/RDS03-094), hDAT/RDS04-010 has many subsegments moving closer to each other on the extracellular side (reddish color in panel **B**) and moving away from each other on the intracellular side (blueish color in panel **C**), thus transitioning to an inward-facing conformation. On panels **D** and **E**, the equilibrated hDAT/RDS04-010 (light blue) and hDAT/RDS03-094 (light pink) complexes are superimposed. On the extracellular side (**D**), TM1e and TM6e and their neighboring subsegments move closer to TM10e, while on the intracellular side (**E**), TM1i swings out towards membrane, with coordinated rearrangement of TMs5i, 6i, and 7i. Panel **F** shows that C𝜶 -C𝜶 distance between Ile67 of TM1i and Leu583 of TM12i (red in panel **G**). Leu583 is relatively stable and selected as a reference for distance measurement. This distance was increased in the hDAT/RDS04-010 simulation not in hDAT/RDS03-094.

The swinging-out of TM1i in the transition towards the inward-facing state, which has been previously observed in the LeuT simulations,^46^ prepares the transporter to release the substrate to the intracellular side. To quantitively evaluate the potential impact of this TM1i rearrangement, we measured the Cα-Cα distance between Ile67 of TM1i and Leu583 of TM12i, and found it fluctuated ∼45 Å in the hDAT/RDS03-094 simulations but increased to ∼55 Å in hDAT/RDS04-010 (**Fig. 4G, F**). This ∼10 Å outward movement of TM1i in hDAT/RDS04-010 was coordinated with the detachment of the N terminal loop from the transmembrane domain, created an opening below the ligand binding pocket to the intracellular water milieu (**movie S1**). Such a process was associated with the tendency of RDS04-010 to be in a more extended conformation (**Fig. 2**). In line with this study, our prior long timescale MD simulations demonstrated that a cocaine analog, CFT, stabilized hDAT in the outward-facing conformation, while an atypical DAT inhibitor, JHW007, steered hDAT towards an inward-facing conformation^38^. However, in that study, the transition to the inward-facing state did not progress far enough to allow for the outward-swinging of the TM1i.

On the extracellular side the ligand binding pocket, the gating residue, Phe320, demonstrated disparate behaviors in the inward-facing and outward-facing conformations. Its χ1 rotamer in the hDAT simulations bound with the sulfide substituted ligands (RDS03-094 and JJC8-089) was populated more in gauche+, while in hDAT/RDS04-010 and hDAT/JJC8-091, this rotamer was in trans and closed the binding pocket from the extracellular side (**Fig. S6**).

In addition, in the simulations of both the hDAT/RDS04010 and hDAT/JJC8-091 conditions, similarly to what has been observed in LeuT^46^, we found that the release of Na2 was associated with outward movement of the TM1i during the transition to the inward-facing conformation.

Taken together, our simulations suggest that ligands with the sulfoxide substitution (RDS04-010 and JJC8-091) induce and stabilize hDAT in an inward-facing conformation, predicting that these compounds would be atypical DAT inhibitors. In contrast, those analogs with the sulfide (RDS03-094 and JJC8-089) prefer an outward-facing conformation and may have a more typical cocaine-like behavioral profile. In the case of JJC8-091, rodent studies support these predictions.^40^ Extensive behavioral investigation of the other analogs is underway.

## Conclusion

This study focused on two pairs of modafinil analogs, JJC8-091 and JJC8-088, and RDS04-010 and RDS03-094, aiming to investigate the impact of the sulfoxide versus sulfide group in affecting conformational changes to hDAT that translates into divergent behavioral profiles. Our QM calculations and conformational search revealed differences in charge distribution and ligand conformation due to these sulfoxide or sulfide moieties. The results of our long MD simulations illustrated that these substitutions prompted the ligand to engage in distinct interactions with hDAT. We found that sulfoxide analogs RDS04-010 and JJC8-091 tend to promote an inward-facing conformation compared to the sulfide analogs of RDS03-094 and JJC8-089. In the transition to inward-facing state, the initial events involved the Na2 escaping from its binding site, followed by the dissociation of the interaction between Asp79 and the pyramidal nitrogen of the ligand. Subsequently, the pyramidal nitrogen to establish an interaction with Asp421. This process is associated with an outward movement of the TM1i that opens the intracellular vestibule to the water milieu.

Thus, the seemingly subtle substituent difference between the sulfoxide versus sulfide in these modafinil analogs strongly affects their preference for either inward-facing or outward-facing DAT conformation. Indeed, it has been found in many instances that similarly minor modifications of small-compound ligands can lead to drastic changes in function at the target proteins. For example, we have reported that introducing an additional methyl group to a synthetic cannabinoid receptor agonist leads to a substantial improvement in its efficacy at the cannabinoid receptor 1.^47^ A comparable subtle alteration in characteristics has also been observed among some 4-phenylpiperazine ligands at the dopamine D3 receptor wherein the (R)-enantiomers are antagonists or weak partial agonists, while the (S)-enantiomers exhibit considerably greater efficacy.^45^

Taken together, our findings of the disparate characteristics of sulfoxide versus sulfide-analogs, along with their atomistic interaction details with hDAT, provide insights into the molecular mechanisms underlying the distinct pharmacological properties of these inhibitors. While our computational approach allows exploration of dynamic behavior and interactions over time, providing insights into the sequence of events in their mechanisms of action, the demanding nature of extensive simulations and analysis required for robust conclusions highlights the non-trivial task of predicting whether a compound functions as a typical or atypical DAT inhibitor. Nonetheless, the gained insights will deepen our mechanistic understanding of the DAT function at atomistic level by fully revealing its functionally relevant conformational spectrum, which will provide the groundwork for the rational design of drugs targeting DAT.

## Methods

### hDAT binding assay

hDAT-HEK293 cell were grown as previously described.^48^ Briefly, HEK293 cells stably expressing human DAT were grown in Dulbecco’s modified Eagle medium (DMEM), supplemented with 5% fetal bovine serum, 5% calf bovine serum, 100 units/mL penicillin/ 100 μg/mL streptomycin and 2 μg/mL puromycin and kept in an incubator at 37 °C and 10% CO_2_. Upon reaching 80−90% confluence, cells were harvested using premixed Earle’s balanced salt solution with 5 mM ethylenediaminetetraacetic acid (EDTA) (Life Technologies) and centrifuged at 3000 rpm for 10 min at 21 °C. The supernatant was removed, and the pellet was resuspended in 10 mL of hypotonic lysis buffer (5 mM MgCl_2_, 5 mM Tris, pH 7.4 at 4 °C) and centrifuged at 14,500 rpm (∼25,000 g) for 30 min at 4 °C. The pellet was then resuspended in binding buffer (50mM Tris, 120 mM NaCl, pH 7.4) Bradford protein assay (Bio-Rad, Hercules, CA) was used to determine the protein concentration and the membranes were diluted to 1000 μg/mL and stored in a −80 °C freezer for later use. Radioligand binding assays were conducted similar to those previously described.^49, 50^ Experiments were conducted in 96-well polypropylene plates containing 50 μL of various concentrations of the test compound, diluted using 30% DMSO vehicle, 300 μL of binding buffer (50 mM Tris, 120 mM NaCl, pH 7.4) 50 μL of [^3^H]WIN35,428^51^ (final concentration 1.5 nM; *K*_d_ = 38.1 nM; NOVANDI Chemistry AB, 78 Ci/mmol SA), and 100 μL of membranes 30 μg/well). All compound dilutions were tested in triplicate and the competition reactions started with the addition of tissue; the plates were incubated for 120 min at 4 °C. Nonspecific binding was determined using 10 μM final concentration of indatraline. Incubations were terminated by rapid filtration through PerkinElmer Uni-Filter-96 GF/C presoaked in 0.2% polyethylenimine, using a Brandel 96-Well Plates Harvester manifold (Brandel Instruments, Gaithersburg, MD). The filters were washed a total of three times with 3 mL (3 × 1 mL/well or 3 × 1 mL/tube) of ice-cold binding buffer. After drying, 65 μL PerkinElmer MicroScint20 Scintillation Cocktail was added to each filter well. Plates were counted using a PerkinElmer MicroBeta Microplate Counter. For each experiment, aliquots of the prepared radioligand solutions were measured to calculate the exact amount of radioactivity added, taking in account the experimentally determined top-counter efficiency for the radioligand. *K*_i_ values have been extrapolated by constraining the bottom of the dose−response curves (= 0% residual specific binding) in the nonlinear regression analysis. *K*_i_ values were calculated using GraphPad Prism 8 version 8.4.0 for Macintosh (GraphPad Software, San Diego, CA) utilizing One site-Fit K_i_ model. *K*_d_ values for the radioligands were determined via separate homologous competitive binding or radioligand binding saturation experiments. *K*_i_ values were determined from at least three independent experiments performed in triplicate and are reported as mean ± SEM.

### Conformational search

The conformational search for each compound was carried out with Conformational Search module implemented in Schrodinger suite (version 2022-3) using OPLS4 force field ^52^ and customized parameters for the missing torsions of RDS03-094 and RDS04-010.

The conformation search was carried out with water as the solvent model, and the search was limited to 1000 steps. During the energy minimization stage, Polak-Robier conjugate gradient (PRCG) was performed, with a cutoff distance of 8.0 Å for van der Waals interactions, 20 Å for electrostatic interactions, and 4 Å for hydrogen bonds. The energy minimization step was performed with 100,000 iterations. The optimal pose for each compound was chosen as the conformation that exhibited the lowest energy after minimization.

### Quantum mechanical calculations

The quantum mechanical (QM) calculations were performed using Jaguar in the Schrödinger suite. The optimized ligand geometries of each compound from the conformational search was used as input in QM calculations. The geometries were first optimized using B3LYP-D3 theory and the 6-31G** basis set in Jaguar. The Poisson Boltzmann Finite element method (PBF) with water as solvent was also applied in the calculation. The same theory and basis set were used for the electrostatic potential calculation and the results were mapped onto the surfaces of constant electron density.

### Optimization of small molecule parameters

To accurately simulate protein ligand interaction without the missing torsions for compounds used in the MD simulation, the ligand parameters were systematically scanning using the force filed builder in Schrödinger suite. The identified missing torsion parameters of RDS03-094, RDS04-010, and JJC8089, were then calculated using the force filed builder with QM calculations.

### Modeling and molecular docking

As RDS04-010, RDS03-094, and JJC8-089 share the same modafinil scaffold as JJC8-091, to establish the hDAT/RDS04-010, hDAT/RDS03-094, and hDAT/JJC8-089 models, we docked the corresponding compound to a well-equilibrated hDAT/JJC8-091 model from previous simulations.^40^ The resulting complex models were then immersed back to original lipid bilayer-water box of hDAT/JJC8-091 to construct corresponding new simulation systems.

### Molecular dynamics simulations

The MD simulations were performed using Desmond MD engine (D. E. Shaw Research, New York, NY) with our previously established protocols for the DAT simulation systems.^38^ OPLS3e force field ^53^ and customized ligands force field parameters were used in MD simulation. Langevin dynamics was performed with NPγT ensemble at constant temperature (310 K) and 1 atm constant pressure with the hybrid Nose-Hoover Langevin piston method ^54^ on an anisotropic flexible periodic cell, and a constant surface tension (x-y plane). The initial complexes systems after docking were first minimized and equilibrated with restraints on the ligand heavy atoms and protein backbone atoms. The restraints were removed during the stage of the production simulations (**Table S1**).

### Structural element definitions and analysis

The structural elements of hDAT were adapted from our previous simulation ^38^ with the following definition: NT (N terminus, residues 58−64), TM1i (the intracellular segment (i) of TM1, residues 65−74), TM1m (the middle segment (m) of TM1, residues 75−82), TM1e (the extracellular segment (e) of TM1, residues 83−92), EL1 (the extracellular loop 1, residues 93−95), TM2e (residues 96−101), TM2m (residues 102−111), TM2i (residues 112−124), IL1 (residues 125−135), TM3i (residues 136−151), TM3m (residues 152−156), TM3e (residues 157−172), EL2 (residues 173−237), TM4e (residues 238−246), TM4i (residues 247−255), IL2 (residues 256−261), TM5i (residues 262−266), TM5m (residues 267−274), TM5e (residues 275−284), EL3 (residues 285−307), TM6e (residues 308−316), TM6m (residues 317−328), TM6i (residues 329−335), IL3 (residues 336−342), TM7i (residues 343−352), TM7m (residues 353−359), TM7e (residues 360−374), EL4a (residues 375−386), EL4b (residues 387−404), TM8e (residues 405−417), TM8m (residues 418−426), TM8i (residues 427−436), IL4 (residues 437−441), TM9i (residues 445−454), TM9e (residues 455−465), EL5 (residues 466−469), TM10e (residues 470−478), TM10m (residues 479−483), TM10i (484−496), IL5 (residues 497−518), TM11i (residues 519−529), TM11e (residues 530−540), EL6 (residues 541−557), TM12e (residues 558−572), TM12i (residues 573−584), and CT (C terminus, residues 585−595). Note that, we redefine the segment range of CT from residues 585−600 to residues 585−595 to avoid the impact of flexible tail of CT in the PIA analyses.

The processing of trajectories and calculation of geometric measurements were performed using a combination of in-house python scripts, MDAnalysis,^55^ and VMD.^56^

The water molecules near the sulfur atoms of the ligands were analyzed using two steps. For each MD frame, the first step identified the water molecules within 3.5 Å of the sulfur atom of the ligand and counted their numbers. The second step calculated the minimum distance between any oxygen atom of the water molecules and the sulfur atom. Then the water counts and the distribution of the minimum distances from the representative MD frame ensemble for each simulated condition are plotted (**Fig. S4B**).

To compute the water counts in the intracellular and extracellular vestibules, the ligands from different conditions were aligned using the same reference, which was based on the stable transmembrane (TM) segments (TM3i, TM3m, TM3e, TM5m, TM5e, TM6e, TM6m, TM6i, TM7m, TM8e, TM8m, and TM8i)^38^. The volume of the extracellular and intracellular vestibules was measured by computing the number of water molecules in each vestibule as described in our previous studies.^38^

The inward-facing ensembles of hDAT/RDS04-010 and hDAT/JJC8-091 were extracted based on the intracellular vestibule water count. Based on the distribution of the intracellular vestibule water counts of the hDAT/RDS03-094 MD frames, we calculated the mean (μ) and one-and-a-half standard deviations (σ) of the water counts, to be the range that defines inward-facing state of each ensemble. A total of ten bootstrapping samples, each containing 1000 frames, were generated for each ensemble for the PIA analyses.

## Author Contribution

K.H.L. and L.S. designed the computational study. K.H.L. carried out computational modeling, simulations, and analysis. G.A. C-H. and A. H. N. designed and carried out the binding assay. All authors took part in interpreting the results. K.H.L. and L.S. prepared and wrote the initial draft, all authors participated in finalizing manuscript.

## Acknowledgements

Support for this research was provided by the National Institute on Drug Abuse–Intramural Research Program, Z1A DA000606 (L.S.) and Z1A DA000389 (A. H. N.). This work utilized the computational resources of the NIH HPC Biowulf cluster (http://hpc.nih.gov).

## Supporting Information

**Table S1.** Summary of MD simulations.

|  | **Runs** | **Length (ns)** |
| --- | --- | --- |
| JJC8-089 | 7 | 19,920 |
| JJC8-091 | 8 | 39,900 |
| RDS03-094 | 7 | 24,120 |
| RDS04-010 | 9 | 40,800 |

**Table S2.** Ligand contact frequencies of the residues interacting with RDS04-010, RDS03-094, JJC8-091, and JJC8-089 that stabilizing inward-facing or outward-facing ensembles. In an MD frame, if the shortest heavy-atom distance between the ligand and any given residue of the protein was within 5 Å, we defined that the ligand forms a contact with this residue. The contact frequency was calculated based on the counting of the contacts per residue and normalized by the ensemble size. The contact frequencies that are above 0.5 for each of the simulated conditions are shaded in cyan. We then calculated the difference between two pairs and highlighted |difference value| ≥ 0.25. Thus, residues with light blue indicate higher contact frequencies with the sulfoxide analogs, and conversely, orange indicates the higher contact frequency with the sulfide analogs.

| Residue | TM | RDS04010 | RDS03094 | JJC8091 | JJC8089 | $\Delta\text{RDS}^*$ | $\Delta\text{JJC}^\#$ |
| --- | --- | --- | --- | --- | --- | --- | --- |
| Phe76 | 1m | 0.83 | 1.00 | 1.00 | 1.00 | -0.17 | 0.00 |
| Ala77 | 1m | 0.57 | 0.86 | 0.82 | 1.00 | -0.29 | -0.18 |
| Asp79 | 1m | 0.73 | 1.00 | 0.87 | 1.00 | -0.27 | -0.13 |
| Ala81 | 1m | 0.35 | 1.00 | 0.31 | 0.98 | -0.65 | -0.67 |
| Val145 | 3i | 0.83 | 0.96 | 0.84 | 0.84 | -0.13 | 0.00 |
| Ile148 | 3i | 0.92 | 1.00 | 0.99 | 0.99 | -0.08 | 0.00 |
| Ser149 | 3i | 0.98 | 1.00 | 1.00 | 1.00 | -0.02 | 0.00 |
| Val152 | 3m | 0.97 | 0.76 | 0.99 | 1.00 | 0.21 | -0.01 |
| Gly153 | 3m | 0.83 | 0.94 | 0.96 | 0.97 | -0.11 | -0.01 |
| Phe154 | 3m | 0.00 | 0.54 | 0.00 | 0.00 | -0.54 | 0.00 |
| Phe155 | 3m | 0.00 | 0.52 | 0.01 | 0.00 | -0.52 | 0.01 |
| Tyr156 | 3m | 0.94 | 1.00 | 1.00 | 1.00 | -0.06 | 0.00 |
| Asn157 | 3e | 0.02 | 0.55 | 0.04 | 0.52 | -0.53 | -0.48 |
| Phe320 | 6m | 0.87 | 1.00 | 0.79 | 1.00 | -0.13 | -0.21 |
| Ser321 | 6m | 0.69 | 1.00 | 0.89 | 1.00 | -0.31 | -0.11 |
| Leu322 | 6m | 0.69 | 0.94 | 0.93 | 0.95 | -0.25 | -0.02 |
| Gly323 | 6m | 0.68 | 0.99 | 0.85 | 0.94 | -0.31 | -0.09 |
| Phe326 | 6m | 0.95 | 1.00 | 1.00 | 1.00 | -0.05 | 0.00 |
| Val328 | 6m | 0.89 | 1.00 | 0.83 | 1.00 | -0.11 | -0.17 |
| Ser422 | 8m | 0.96 | 1.00 | 1.00 | 1.00 | -0.04 | 0.00 |
| Ala423 | 8m | 0.81 | 0.97 | 0.96 | 0.94 | -0.16 | 0.02 |
| Gly425 | 8m | 0.85 | 0.56 | 0.89 | 0.58 | 0.29 | 0.31 |
| Gly426 | 8m | 0.91 | 0.89 | 0.99 | 1.00 | 0.02 | -0.01 |
| Ser429 | 8i | 0.80 | 0.53 | 0.41 | 0.19 | 0.27 | 0.22 |
| Ala479 | 10m | 0.37 | 0.67 | 0.12 | 0.56 | -0.30 | -0.44 |
| Ile484 | 10i | 0.84 | 0.87 | 0.74 | 0.91 | -0.03 | -0.17 |

\* $\Delta\text{RDS}$ is the difference using RDS04010 - RDS03094, # $\Delta\text{JJC}$ is the difference using JJC8091- JJC8089.

**Figure S1.**
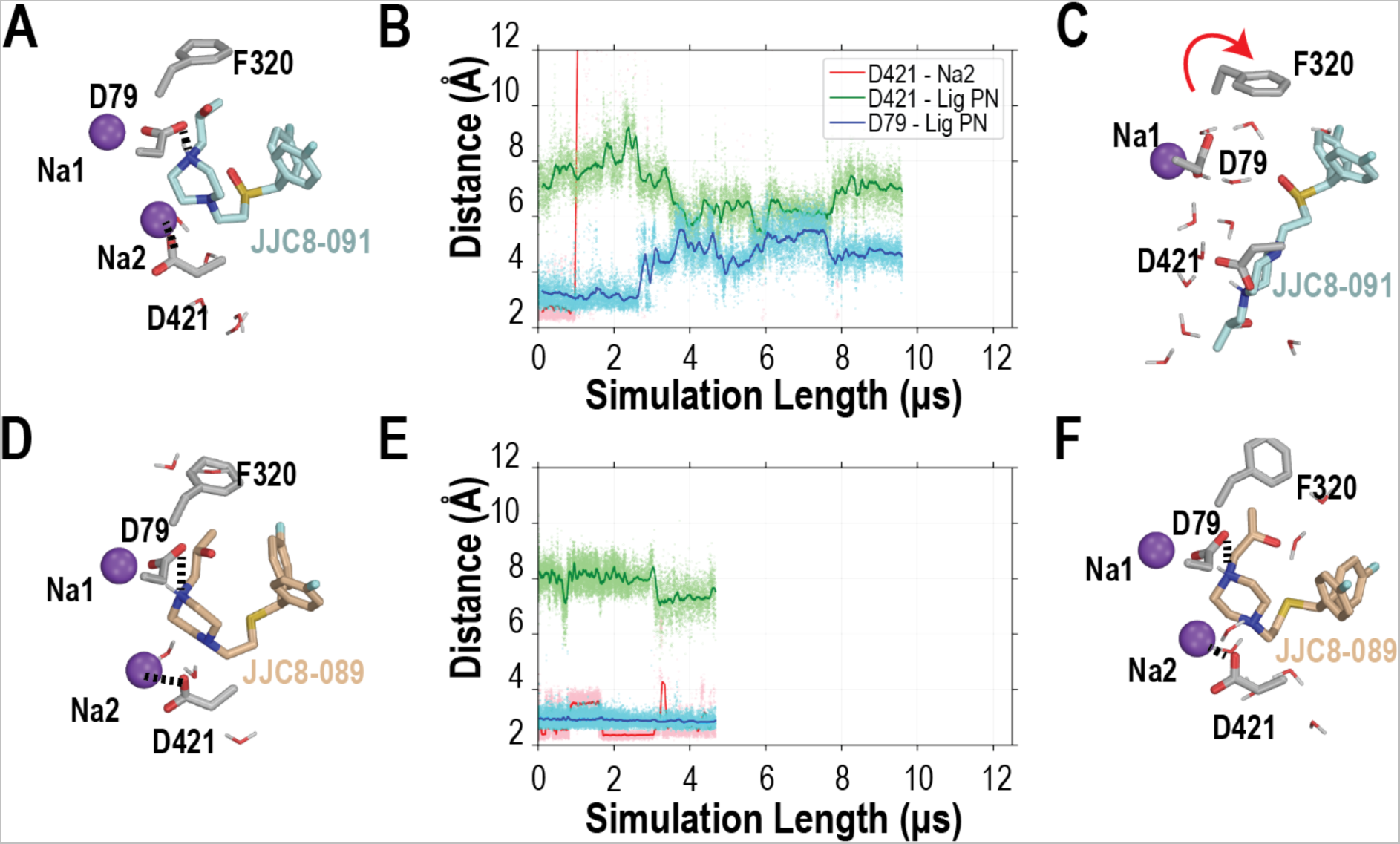
The bound of JJC8-091 cannot stably interact with Asp79 resulting in the dissociation of Na2. The evolution of the distances between the pyramidal N of the ligand and these two residues are shown in panel **B** for JJC8-091 and panel **E** for JJC8-089. The beginning binding poses of JJC8-091 and JJC8-089 are shown in **Fig. S2A** and **D** respectively, and the representative snapshots of the binding poses of JJC8-091 and JJC8-089 are shown in **Fig. S2C** and **F** respectively.

**Figure S2.**
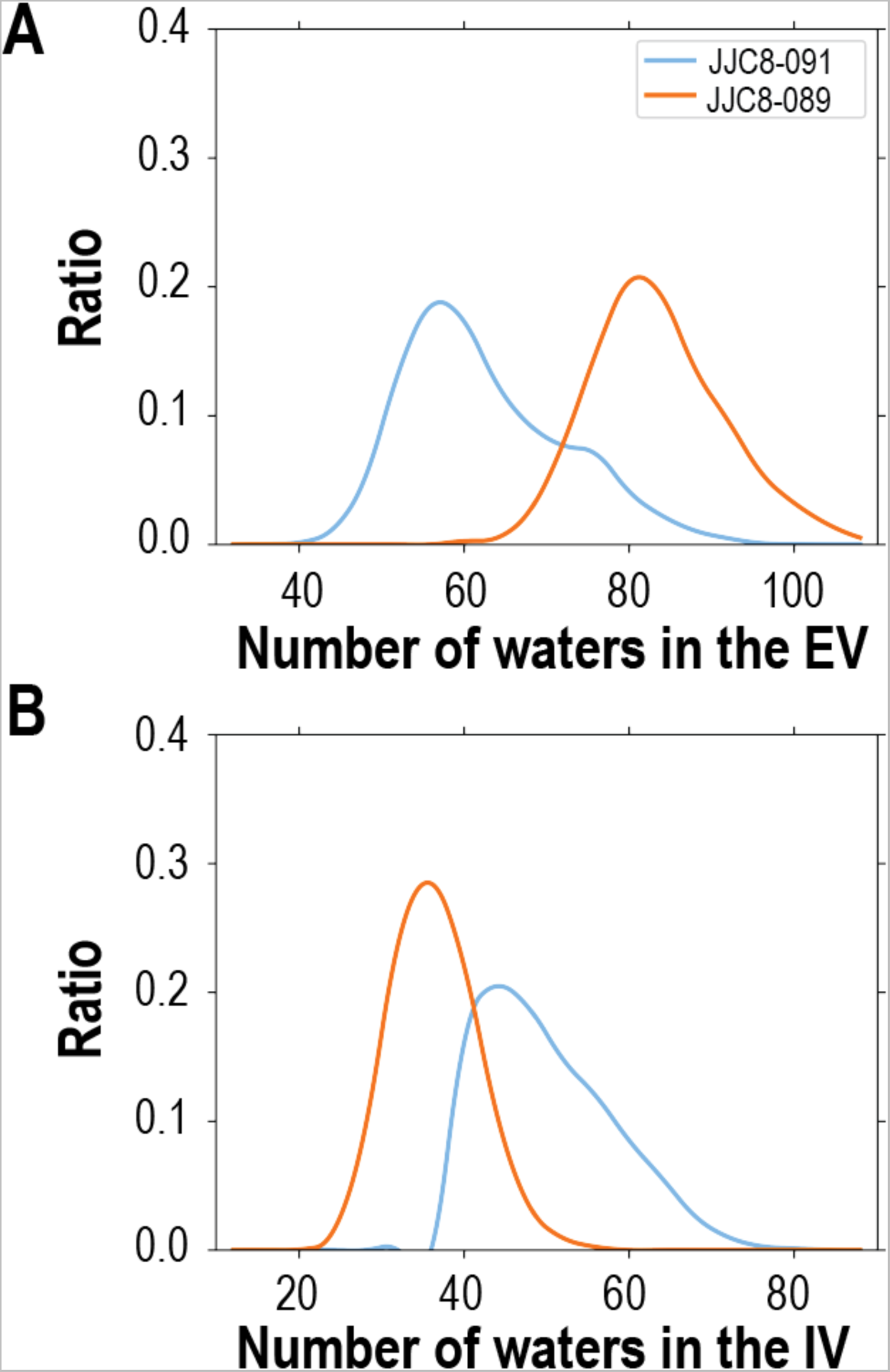
JJC8-091 and JJC8-089 stabilized DAT in inward-occluded ad outward-open conformations, respectively. We measured the volumes by counting the water molecules in the vestibules (see Methods) and found that hDAT/JJC8-089 (orange) has ∼20.3 more waters in the extracellular vestibule (**A**) and ∼6.4 fewer waters in the intracellular vestibule (**B**) than hDAT/JJC8-091 (blue).

**Figure S3.**
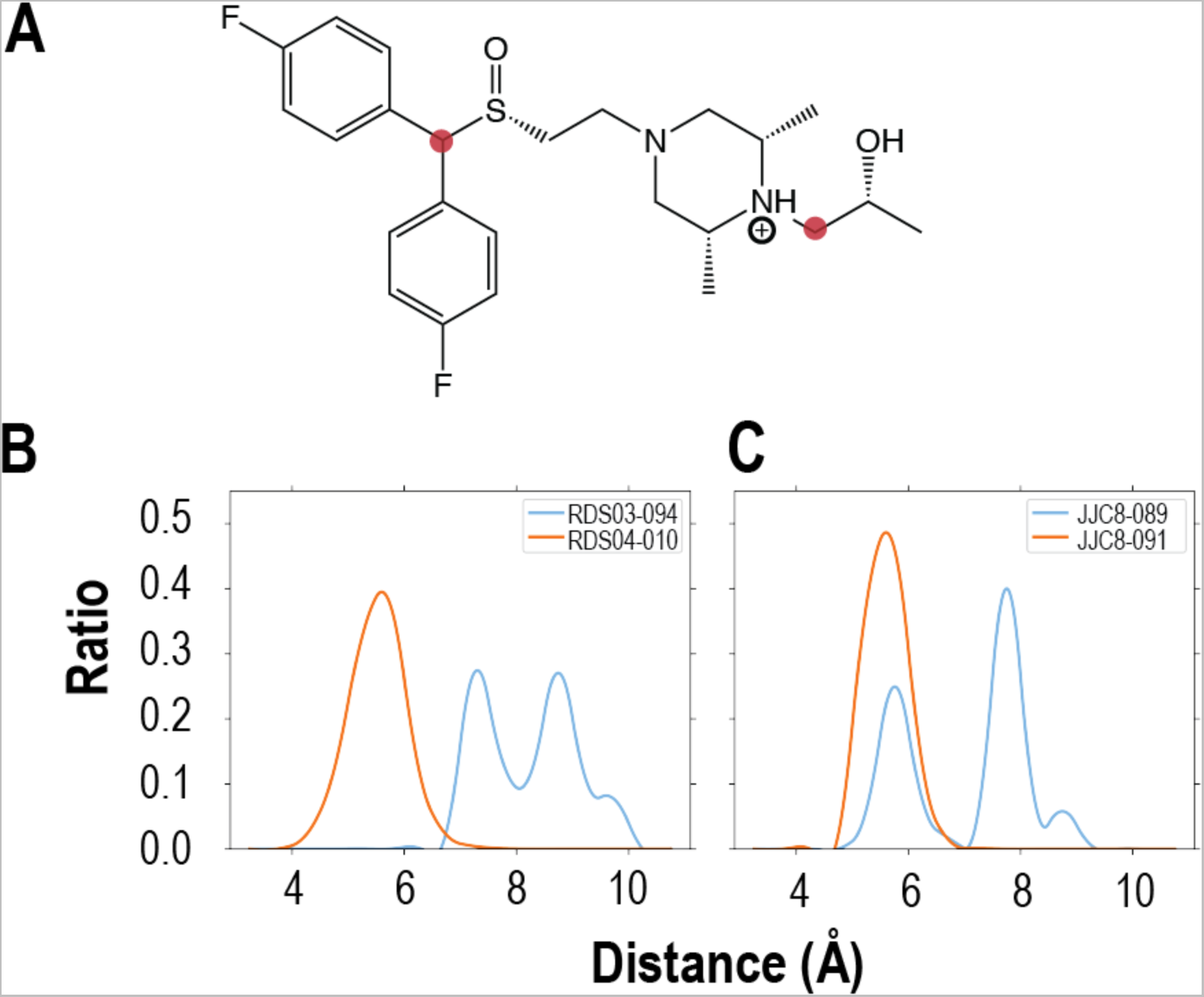
Distribution of ligand end to end distance. Ligand end-to-end distance was calculated using two defined carbon atoms as shown in red dots (**A**). Distribution of ligand end-to-end distance of those carbons of hDAT/RDS04-010 and hDAT/RDS03-094 is shown in (**B**), and of hDAT/JJC8-091 and hDAT/JJC8-089 is shown in (**C**).

**Figure S4.**
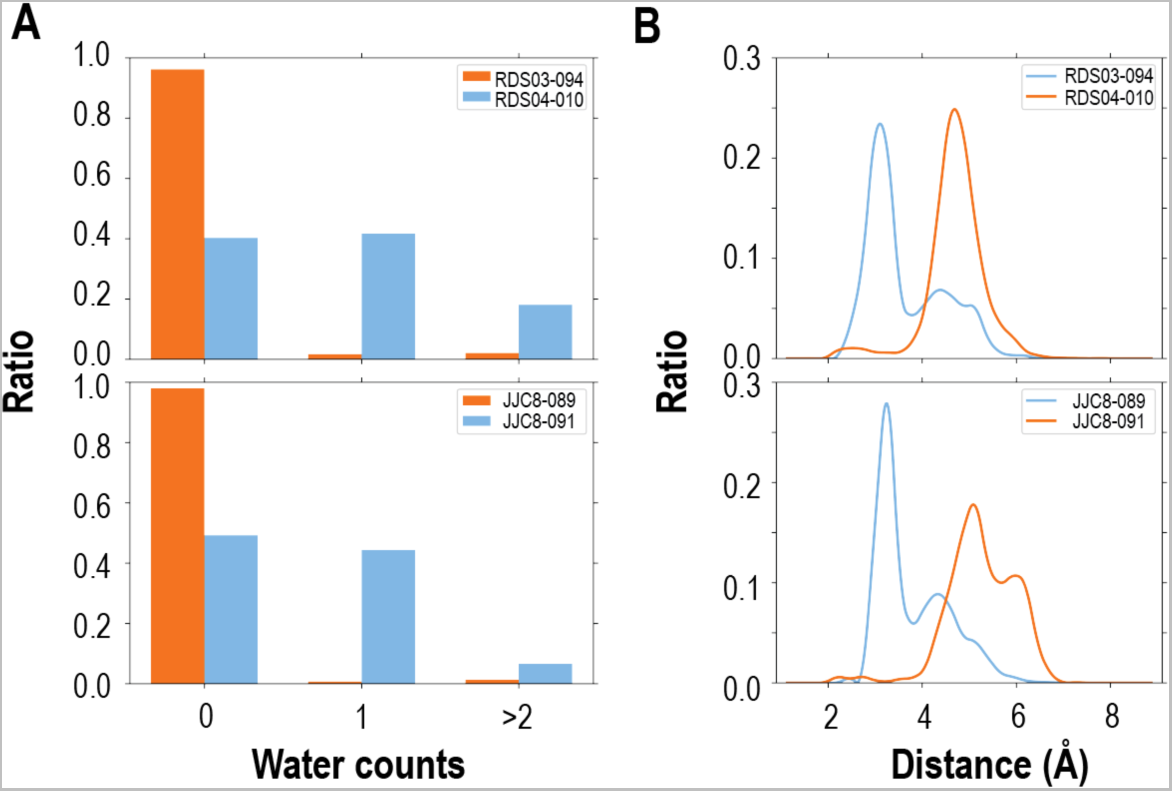
The sulfoxide substitution attracts more water than sulfide. The difference between inward-facing and outward-facing ensembles was shown in terms of the number of water counts around the sulfur atom of substitution sulfoxide and sulfide (**A**) and the distribution of minimum water-sulfur atom distance (**B**). Upon counting the water molecules near the sulfur atom (within 3.5 Å), it was found that there are more water molecules near sulfoxide compared to sulfide (panel **A**). For substitution of sulfoxide, it’s 59.8% for RDS04-010 and 50.9% for JJC8-091 that having more than one water molecule. For substitution of sulfide, it’s 3.8% for RDS03-094 and 2.0% for JJC8-089. In panel B, the minimum distance between the closest water to the sulfur atom is used to compare the water nearby sulfoxide and sulfide substitution. Those water nearby sulfoxide can form a single peak around 3.1 Å and 3.3 Å for both RDS04-010 and JJC8-091, suggesting a direct hydrogen bond was formed between sulfoxide and water. For sulfide substitution, the peaks are around 4.7 Å and 5.1 Å for RDS03-094 and JJC8-089 respectively.

**Figure S5.**
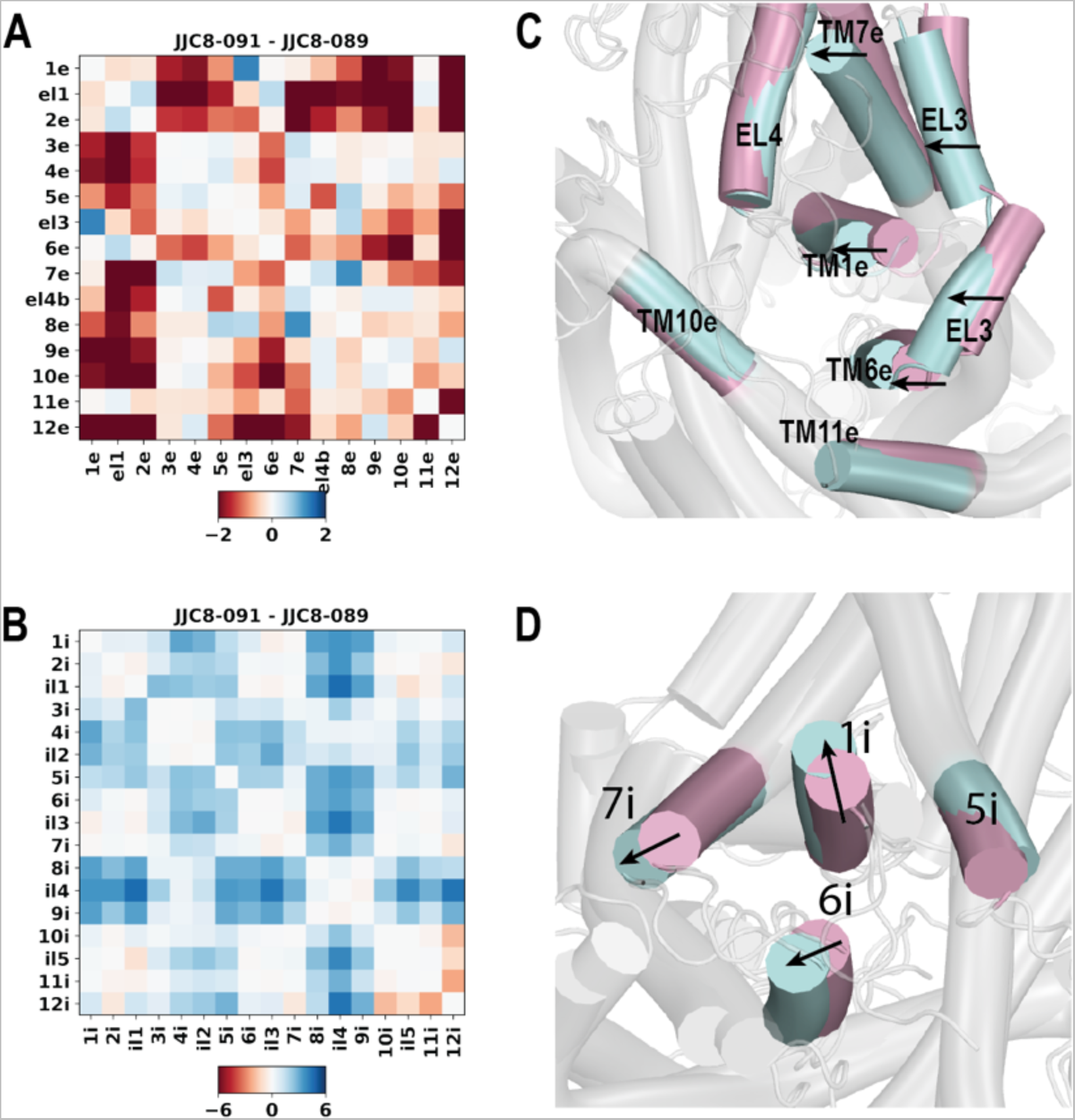
hDAT/JJC8-091 has a tendence to be more inward-facing compared to hDAT/JJC8-089. Extracellular (**A**) and intracellular (**B**) PIA distance matrixes were shown more inward-facing (blueish color in panel **B**) for JJC8-091 and more outward-facing (reddish color in panel **A**) for JJC8-089. The superimposition of the equilibrated DAT/ JJC8-091(light blue) and DAT/ JJC8-089 (light pink) complexes show the inward movement of TM1e, EL3, and TM6e (**C**) and the outward movement of TM1i (**D**).

**Figure S6.**
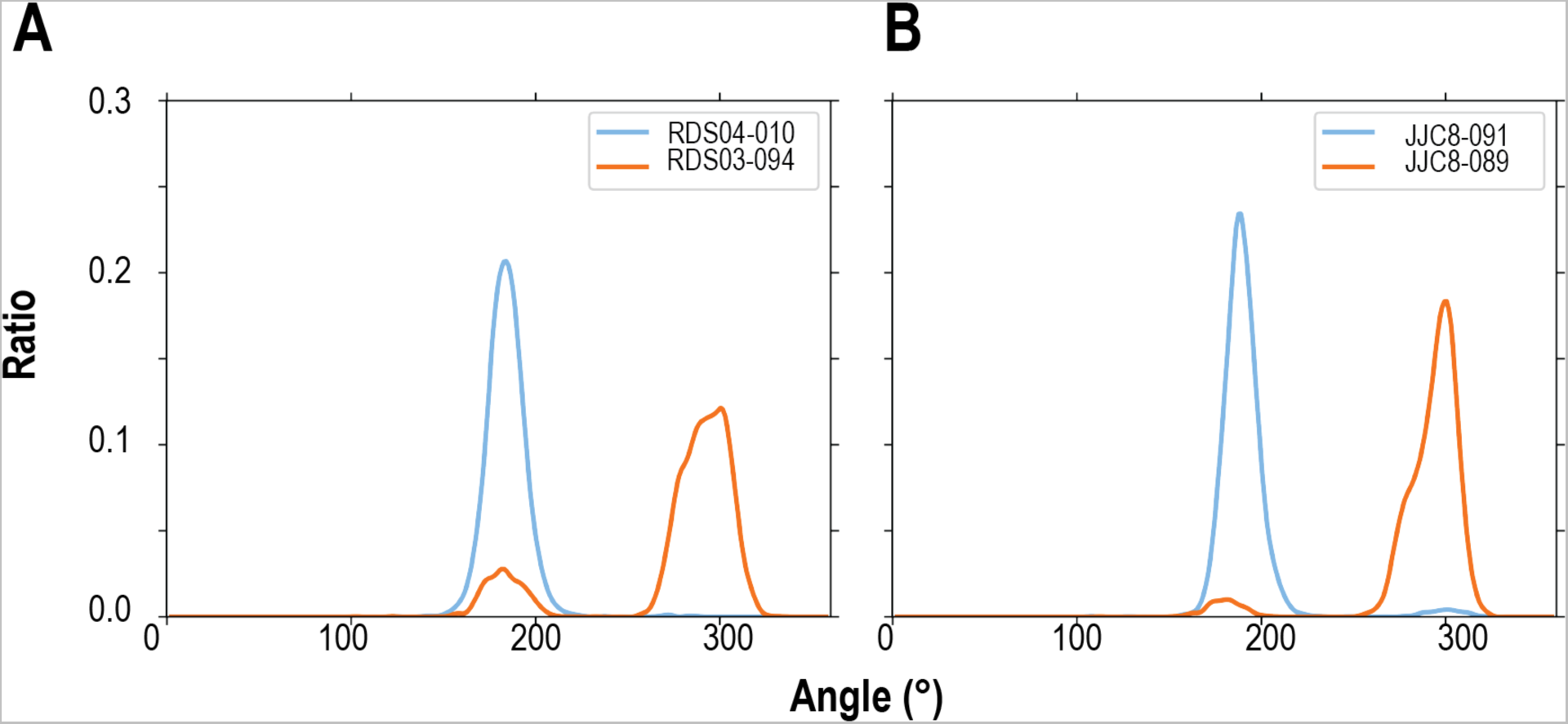
F320 χ1 sampled more trans conformation for compounds with substitution of sulfoxide. The distribution of F320 χ1 angle for RDS04-010 and RDS03-094 (**A**), and for JJC8-091 and JJC8-089 (**B**).

**Movie S1. The transition to an inward-facing conformation observed in a representative hDAT/RDS04-010 MD simulation trajectory.**

The intracellular portion of the TM1 (TM1i) and N-terminal loop are colored in magenta. Na ions are shown in purple spheres. The carbon atoms the bound RDS04-010 are in green. The dissociation of the Na2 leads to the swinging out the TM1 and detachment of the N-terminal loop from the TM domain, while RDS04-010 transitions to a more extended conformation in the newly created cavity by these rearrangements.

